# Using General Algebra to Model the Directed Evolution of an Asexual Population

**DOI:** 10.1101/2025.11.22.689897

**Authors:** Kangbien Park

## Abstract

Humans have long employed directed evolution (DE) to engineer desired biological traits. In this paper, I introduce an algebraic framework that provides a quantitative representation of the general phenotypic traits of asexual populations, enabling the systematic modeling of DE processes. Within this framework, key evolutionary quantities such as the time required for a DE to reach a desired trait or the probability of arriving at a target algebra can be computed or qualitatively analyzed using principles from the evolutionary dynamics of an asexual population. As illustrative examples of trait-representing algebras, I evolve an integer, a two-dimensional vector, a four-dimensional vector with modulo-4 elements, a 2 by 2 matrix, and a cyclic algebra. The generations needed to reach the objective algebra in the DE simulations were consistent with those predicted by the theoretical analysis. Furthermore, I propose a method for mathematically designing evolutionary pathways that minimize the generations needed to reach the desired algebra, offering a key criterion for improving the efficiency of DE. Finally, I discuss how this algebraic approach can be applied in practical experimental setups and outline directions for future research in algebraic modeling of evolution.

## 2 Introduction

Directed evolution (DE) has long been of interest to humans. Since prehistoric times, humans have shaped the evolutionary trajectories of crops and livestock through artificial selection. With the advent of modern scientific techniques, DE of asexual microbial populations has become possible, enabling the evolution of traits of interest Wright et al. 2019, Arias-Sánchez et al. 2024. Despite the ongoing success of DE in asexual populations using qualitative chemical and biological methods, achieving more efficient and precise control of DE requires a well-organized quantitative approach Tack et al. [2020].

For example, if experimenters aim to evolve *E. coli* to produce a novel medicine, they could design schemes based on qualitative biological and chemical methods Lee et al. [2011]. However, such approaches have fundamental limitations without mathematical analysis, as there is no precise way to predict the duration of the process, the probability of success, or even whether the qualitative scheme will be effective given the inherent stochasticity of evolution Alexander [2023]. Furthermore, even if more efficient evolutionary pathways exist—taking advantage of the complex dynamics of existing mutations—these paths may remain undiscovered without quantitative guidance. As the trait of interest becomes more complex, this limitation is exacerbated, since qualitative reasoning alone cannot rigorously track the evolution process. By analogy, one might fly a hot-air balloon without extensive mathematics to reach moderate heights, but no spacecraft could be designed without the rigorous application of mathematically formulated aerodynamics.

Fortunately, with the continuous development of evolutionary dynamics and population genetics, applying quantitative methods to the directed evolution (DE) of asexual populations has become increasingly promising Desai et al. [2007], Corrao et al. [2024]. In addition, rapid advances in computer science and artificial intelligence now allow numerical simulations with large datasets like never before, providing powerful tools to design evolutionary pathways from mathematical and physical perspectives. Despite these opportunities, quantitative control of asexual population evolution remains a nascent field that requires substantial theoretical development before practical implementation Charlebois [2023]. Achieving this goal greatly benefits from tools that can intuitively represent traits of interest as mathematical objects, because they allow precise definition of corresponding fitness landscapes and mutations, thereby enabling rigorous calculation of diverse quantities involved in DE. In this paper, I introduce a framework for defining the most general form of algebra to represent the evolution of any phenotypic trait, including traits encoded by genetic sequences, and demonstrate how it can be applied to model the DE process. Although the model does not rely heavily on detailed knowledge of molecular biology, the model still remains promising because it provides a quantitative, phenotypic representation of evolution—essentially formalizing the processes that humans have been shaping for tens of thousands of years, long before the discovery of genetic material. As examples of applying the algebraic model, I evolve various types of algebras—including integers, vectors, matrices, and finite cyclic algebras—and theoretically analyze the expected progression of their evolution.

## 3 Theory and Methodology

The main objective of this section is to introduce a mathematical framework for modeling the directed evolution (DE) of an asexual population and to present the conditions required or advisable for effective DE.

1. Define a suitable algebra *A* that represents the trait(s) of interest *T*_1_, *T*_2_, …, *T*_*n*_ with an operation ∗, so that *A* = (*A*(*T*_1_, *T*_2_, …, *T*_*n*_), ∗). Here, the trait(s) can be any quantifiable characteristic of a biological entity, including DNA sequences. The set *A* can be either finite or countably infinite: for a finite set of *k* possible mutations, *A*_finite_ = {*A*_1_, *A*_2_, …, *A*_*k*_ }; for an infinite set, *A*_infinite_ = {*A*_1_, *A*_2_, …, *A*_*k*_,}. Furthermore, when a new mutation occurs, the algebraic element corresponding to the resulting mutation is determined according to the rules defined by the operation ∗.
2. Define a *potential* Φ such that

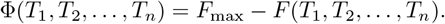

where *F* (*T*_1_, *T*_2_, …, *T*_*n*_) is a fitness landscape in *n* dimensions determined from the *n* independent traits *T*_1_, *T*_2_,, *T*_*n*_ that define the algebra.
3. Define a function *f*: *A* → ℝ that assigns a selection coefficient Φ = *f* (*x*) to each element *x* ∈ *A*.

With such a well-defined algebra *A* = (*A*(*T*_1_, *T*_2_, …, *T*_*n*_), ∗) and function *f*: *A* → Φ, in a finite number of generations, any algebra *A* can evolve to the objective algebra under the following conditions.

1. The degrees of freedom of mutation are limited; that is, mutations can only—or *effectively* only—occur within the algebraic set(s) of interest.
2. The population size is sufficiently large so that genetic drift is negligible and deterministic behavior dominates; that is, *N* ·*s* ≫ 1 and *s* ≪ 1.
3. Evolution proceeds in the direction of decreasing potential when

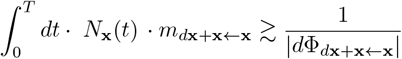

for some unit generation interval *T*.
4. The range of algebras from the initial algebra to the target algebra should be *explorable* through mutation. Here, the definition of *explorability* includes the possibility of establishing a population. In other words, even if a point is reachable by mutation, it is not considered explorable if a population cannot become established there.
5. (*Weak condition*) Clonal interference is useful for sophisticated control.

To illustrate the above definitions and conditions, a one-dimensional integer algebra (*A*_integer_, +), where mutation occurs in the +1 or − 1 direction, and a two-dimensional vector algebra (*A*_1*×*2_, +), where mutation occurs only in the +(1, 0) or +(0, 1) directions are used.

Let us assume that the shape of the fitness landscape for (*A*_integer_, +) is shown in Fig 1 (a), and the corresponding potential is illustrated in Fig 1 (b). Note that Δ*s* = − ΔΦ. The potential is defined to intuitively represent the idea that the state of an algebraic system, depicted as a red ball, *always* slides down toward a more locally stable position—analogous to an object *deterministically* rolling down to the local minimum (stable position) of a hill. For such a deterministic evolution to occur, Condition 1 and Condition 2 must both hold.

**Figure 1.**
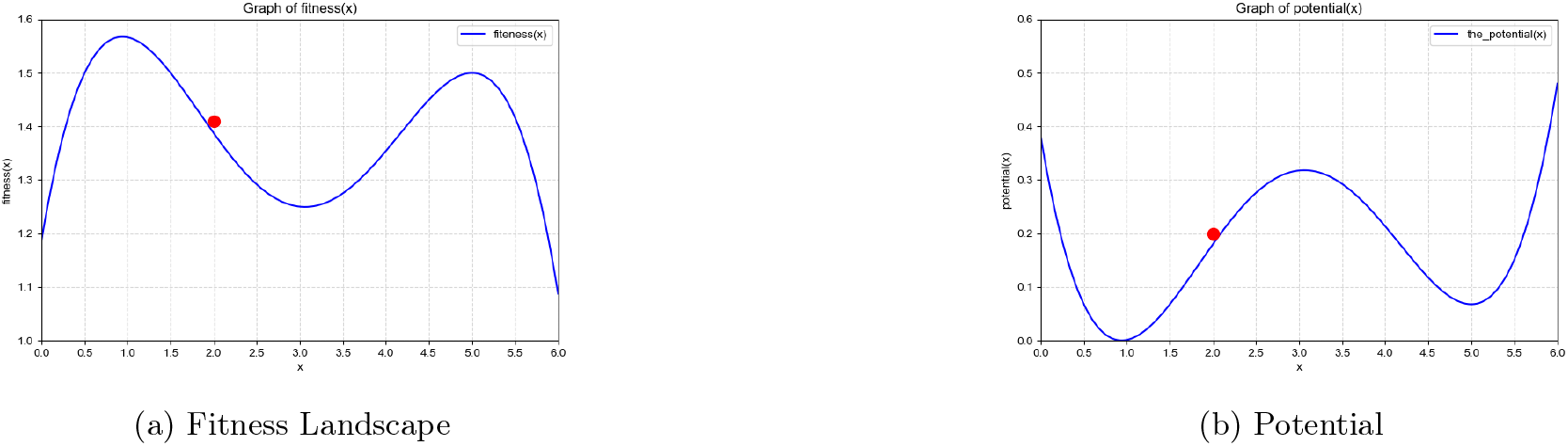
Fitness landscape and corresponding potential.

If Condition 1 is not satisfied, the potential cannot be properly defined within the limited information provided solely by the algebraic set (*A*_integer_, +). Here, “effectively” means that, within the local generational range of interest, the potential governing the trait of interest can be regarded as time-independent, while mutations occur only in a limited manner consistent with the elements of the algebraic set. Moreover, if Condition 2 is not satisfied, drift plays a large role in the dynamics, and the deterministic analysis may no longer be valid.

To explain Condition 3, refer to Fig 2 (a), which illustrates the accessible states after mutation. Each new algebra corresponding to a mutation is obtained by adding or subtracting 0.5 from the original algebra. Since the probability of establishment for any mutation is given by *P*_*establishment*_ ∝ Δ*s* = − ΔΦ_*±*0.5+2←2_, we find that only the mutation in the leftward direction can become established, because ΔΦ *<* 0 only in this case, resulting in a positive establishment probability.

**Figure 2.**
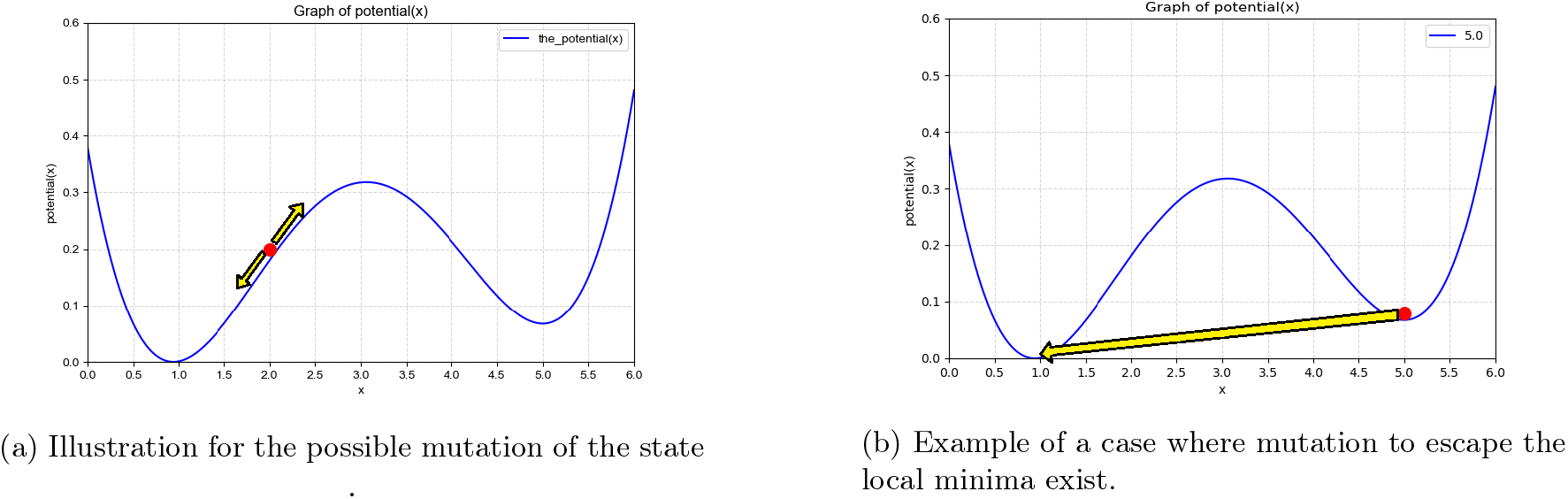
Examples of the evolutionary process in one dimensional algebra set.

Considering the entire population with trait *x* = 2 as *N*_*x*=2_(*t*), the probability of mutation establishment per generation is

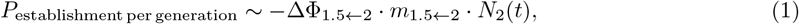

and over a generational interval *t*,

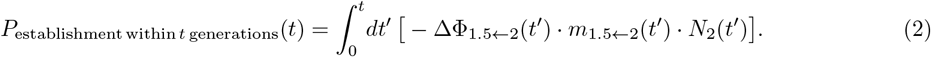

From these relations, the timescale for deterministic behavior can be estimated from the condition *P*_establishment_ ~ 1 which implies the Condition 3. Therefore, if Condition 3 holds, establishment always occurs, and the population evolves deterministically downward along the potential landscape. Furthermore, even if the system resides in a local minimum, as shown in Fig 2 (b), the state can still reach the global minimum if a mutation pathway such as *x* = 5 → *x* = 1 exists, that is, if *m*_1←5_ ≠0.

To understand Condition 4, refer to Fig 3, which illustrates the evolution of a two-dimensional (2-D) vector. Starting from Fig 3 (a), observe that mutation toward the (0, 1) position decreases the potential, whereas mutation in the (1, 0) direction does not provide such an advantage. Consequently, the state cannot move in the *x* direction, and thus (1, 0) is not explorable by definition. Although this qualitative intuition holds for the case shown in Fig 3 (a), a more general treatment of various potential forms requires quantitative analysis. For this purpose, consider Fig 3 (b), where the potential difference is given as 0.5*s* in the (1, 0) direction and 0.8*s* in the (0, 1) direction. Let us now assume that the evolutionary reaction from (0, 0) to (2, 0) is of interest. Here, I define the term *evolutionary reaction* as the DE process in which the objective algebra is established from the initial algebra.

**Figure 3.**
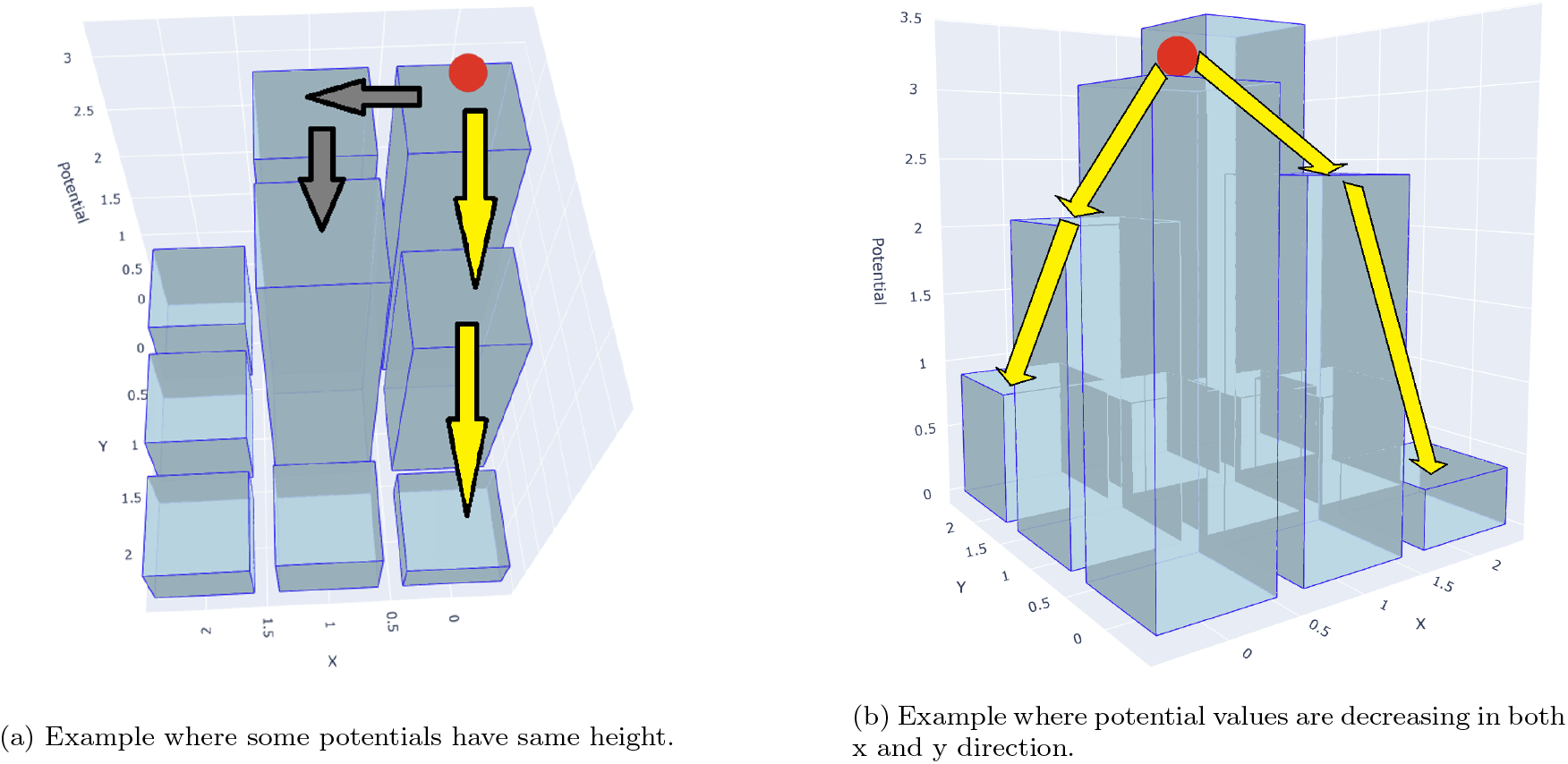
Examples of potential for 2-D vector. To get a precise value for ΔΦ, multiply a selection coefficient *s* to the z values. For (a), ΔΦ_(1,0)←(0,0)_ = 0, ΔΦ_(0,1)←(0,0)_ = −0.8 · *s*, ΔΦ_(0,2)←(0,0)_ = −1.6 · *s*. For (b), ΔΦ_(1,0)←(0,0)_ = −0.5 · *s*, ΔΦ_(0,1)←(0,0)_ = −0.8 · *s*, ΔΦ_(2,0)←(1,0)_ = −2 · *s*, ΔΦ_(0,2)←(0,1)_ = −1.2 · *s*.

Returning to the potential in Fig 3 (b), even though the *y*-direction has a larger potential difference than the *x*-direction, the (0, 1) mutation can still become established before the (1, 0) mutation is fixed. This implies that the final evolutionary outcome can still reach (2, 0), even though the potential of the intermediate state (1, 0) is relatively weak under selection. In this case, for the potential of Fig 3 (b), the positions (0, 0), (1, 0), (2, 0), (0, 1), and (0, 2) are all *explorable*.

To estimate the probability that (2, 0) is eventually established, note that the time to establishment can be expressed as

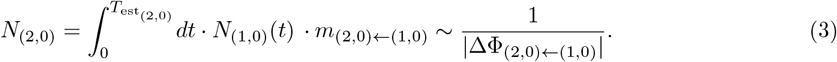

Moreover, from the traveling-wave model Fisher [2013], Desai et al. [2007], the population growth of the leading edge (or “nose”) of the mutation distribution is described as

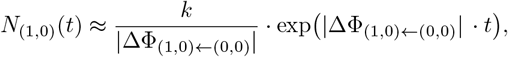

where *k* is a constant used to account for empirical evolutionary effects within a given environment and parameter set, relative to theoretical expectations. In this study, *k* is approximated as *k* ≈ 1, following the theoretical analysis of Desai et al. [2007].

Now, substituting *N*_(1,0)_(*t*) into Eq (3) gives

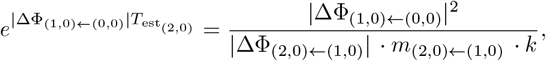

which yields

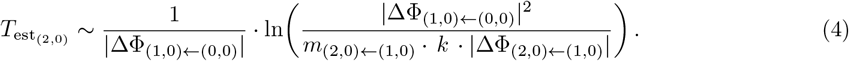

It is also straightforward to estimate the time required for the establishment of (1, 0) by noting that

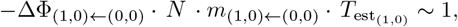

which gives

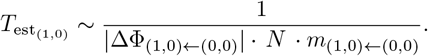

The same formulation applies to evolution in the *y* direction, yielding

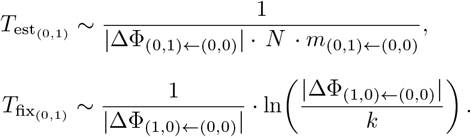

Therefore, the probability

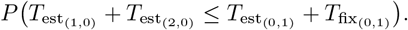

is of interest. Although a more detailed quantitative treatment using generating functions could yield an exact distribution, here, only a qualitative interpretation based on the inequality (5) is provided.

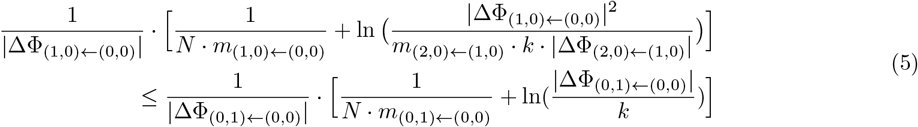

In this relation, the left-hand side term inside the logarithm can be rewritten as

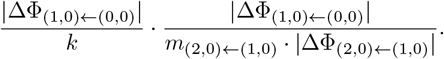

Then, even if 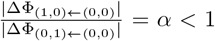. the inequality (5) can still hold when *m*_(2,0)←(1,0)_ and |ΔΦ_(2,0)←(1,0)_ | are sufficiently large and *α* is not too small. This observation implies the occurrence of *relative tunneling*, a phenomenon in which states explore potentials that do not have the lowest values among available mutations. Such relative tunneling, in which intermediate traits are established but do not eventually fix, can play a critical role, as these intermediate traits may serve as essential steps guiding the evolution toward the objective algebra.

Lastly, Condition 5 follows from the qualitative observation that intensive clonal interference (CI) Park and Krug [2007], De Visser and Rozen [2006] enables existing mutations to explore a broader range of possibilities. Notably, this observation aligns with the result of inequality (5), which indicates that a higher mutation rate leads to more explorable points via relative tunneling. Furthermore, because evolutionary speed is reduced under CI compared to successive sweep scenarios Desai et al. [2007], Kuo et al. [2025], it becomes more effective for experimenters to intervene and control the evolutionary process at appropriate times.

Although there is no strict causal relationship, the combination of slower evolutionary speed due to interference and stronger CI enhancing the number of explorable algebra elements provides the following intuition: similar to how precise steering of a car requires slower speed, the reduction of evolutionary speed from CI allows for improved sophistication in directing evolution. Importantly, despite CI slowing the evolutionary process while expanding explorability, the overall time to reach the objective can still be reduced, as illustrated in the matrix evolution examples below. Therefore, to increase the success rate in directing evolution—particularly when relative tunneling is relevant—conditions satisfying 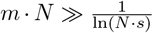 Desai and Fisher [2007] that ensure intensive clonal interference are particularly advantageous. This phenomenon can be aptly summarized by the proverb: *“A stitch in time saves nine*.*”*

In summary, with the defined mathematical system and the specified conditions, the goal of directed evolution using algebraic modeling can be intuitively illustrated by Fig 4, where the algebra considered is a 2-D vector with continuous elements. Based on the potential shown in Fig 4, the vector is likely to evolve toward the target position (12, − 7) via mutation, provided that Condition 3 is satisfied. It is important to note that Condition 1 restricts mutations to occur only within the plane, prohibiting any movement along the *z*-direction or any higher-dimensional axes. Consequently, the task in directed evolution is to carefully design the potential so that the current state successfully reaches the target position.

**Figure 4.**
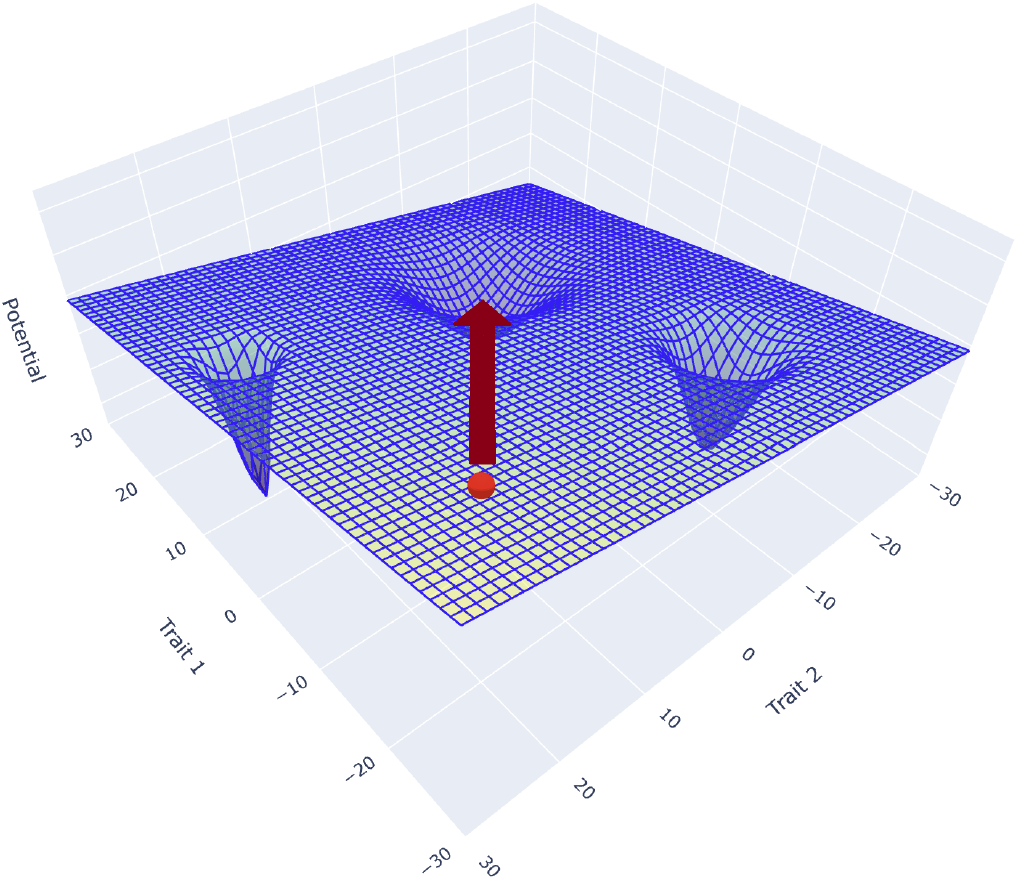
Intuitive Representation of the Objective of Directed Evolution. The objective of directed evolution is intuitively represented using a 2D vector evolution scheme. The essential idea is to manipulate the potential and the mutation rate so that the population, through evolutionary dynamics, progresses toward the objective vector.

The following are possible examples of diverse algebraic modeling scenarios in which the evolutionary parameters satisfy the five conditions described above. Here, assume that the potential is either monotonically decreasing or locally convex in an *n*-dimensional space, with the objective algebra corresponding to the global minimum. In cases where the potential is locally convex, it can be approximated by a quadratic convex function through a series expansion. Accordingly, the potentials for the examples below are designed to be linear or quadratic from the outset, except in the case of cyclic evolution.

1. Evolution of integers, where the mutation law is defined as follows: addition of 1 with probability 0.6, addition of 3 with probability 0.2, or multiplication by 6 with probability 0.2 applied to the original element. The objective number to reach is any random number, for example, 1346, and the potential is defined as

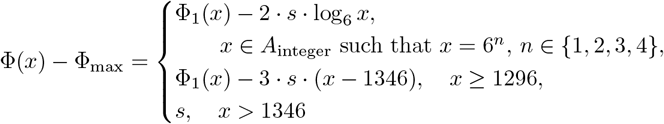

where

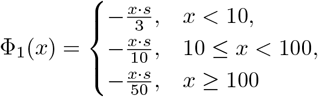 For simulation, the selection coefficient is set to *s* = 0.02. The corresponding figure of the potential is provided in the appendix **??**.
2. Evolution of a 4-D vector (*a, b, c, d*), where each component is a modulo 4 integer, and the mutation occurs by adding 1 to one of the four components with probability 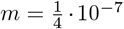. Two evolutionary scenarios were considered: one from (0, 0, 0, 0) to (2, 3, 1, 2), and the other from (0, 0, 0, 0) to (1, 1, 1, 1). The potential for the first case is

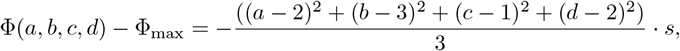

while the potential for the second case is

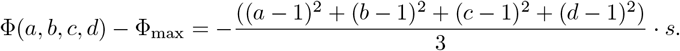 For simulation, the selection coefficient is set to *s* = 0.01 for the (2, 3, 1, 2) case and *s* = 0.03 for the (1, 1, 1, 1) case.
3. Evolution of a 2×2 matrix starting from 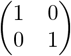, where the determinant and trace govern the evolution of the matrix. Mutation occurs by adding a matrix 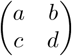 selected from 24 different combinations with *a, b, c, d* ∈ {0, ± 1, ± 2, 3 }, each occurring with probability 1*/*24. The objective determinant is 3 and the trace is 8, so the potential is defined as

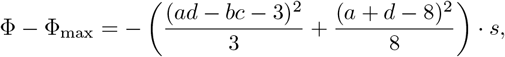

with four independent variables *a, b, c, d*. The specific forms of the mutation matrices are provided in the appendix. For simulation, the selection coefficient is set to *s* = 0.01.
4. Finitely bound cyclic evolution (Rock-Paper-Scissors type evolution). When the algebra is cyclic with a finite set size, one can consider cyclic evolution in which the trait ultimately returns to itself. In this case, the potential is defined based on the proportion of specific mutations. For simulation, we used three mutations: Rock, Paper, and Scissors. The potential for Rock is given by

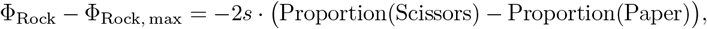

with analogous definitions for Paper and Scissors. For simulation, the selection coefficient is set to *s* = 0.01.

Lastly, simulations are performed using the operator model Park and Bae [2024], which efficiently represents the DE process of the asexual population. Moreover, ChatGPT was used to generate parts of the plotting code and to assist in designing and debugging specific computational components.

## 4 Results and discussion

Fig [5] shows the results for the *N* = 10^7^ 2-D vector evolving under the potential of Fig 3 (b), which requires relative tunneling to reach the objective algebra (2, 0). As predicted by inequality (5), when the mutation rate is sufficiently high, around *m* ~ 5 · 10^−6^, the success rate approaches 1. Interestingly, the success rate is lowest when *m* ~ 1 · 10^−7^: although many mutations occur to explore both (1, 0) and (0, 1), there are not enough mutations to reach (2, 0).

**Figure 5.**
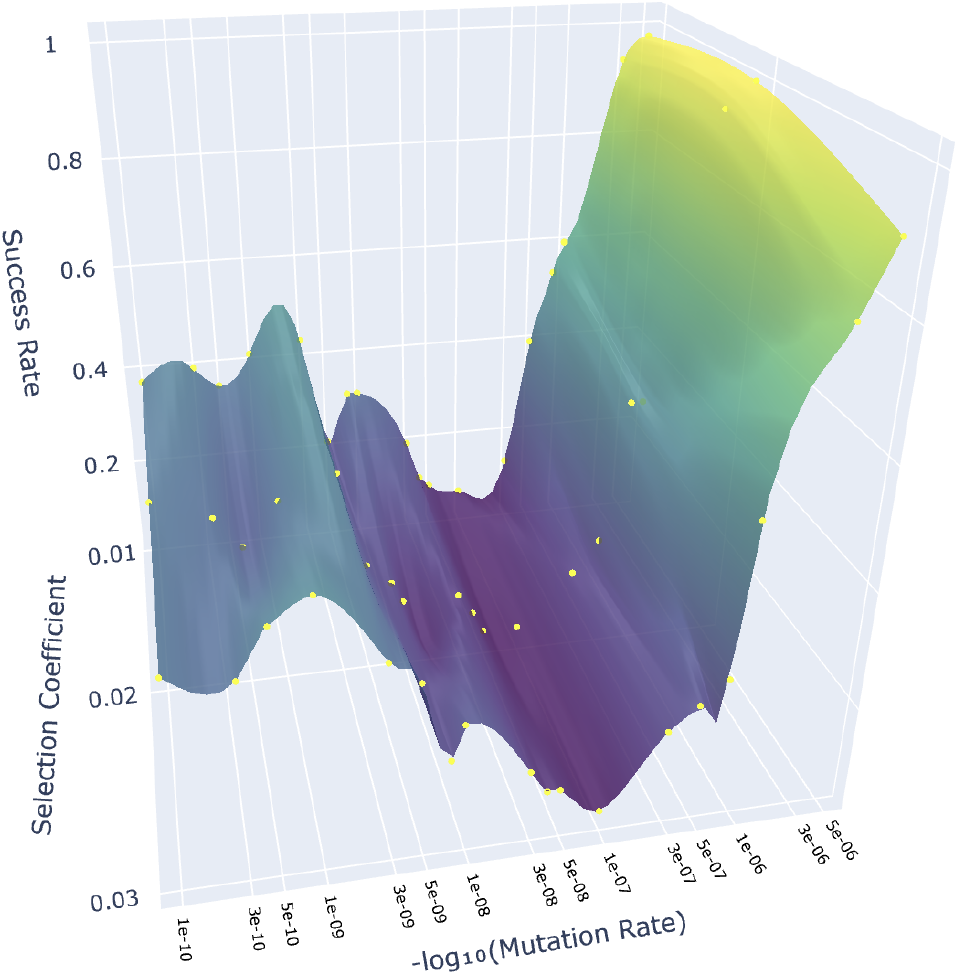
Success rate of 2-D vector evolution measured from diverse selection coefficient and mutation rate values. The yellow dots indicate the success rate values obtained from simulations for the specified parameter combinations, and the remainder of the surface is interpolated from these sampled points.

Notably, for *m* ~ 10^−9^, the success rate is approximately 0.5. At this mutation rate, mutations occur just enough to allow the bare establishment of either (1, 0) or (0, 1) vectors, so the outcome depends solely on whether the state “drops” to the left or right, giving a probability of 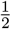.

Finally, when *m <* 10^−9^, the success rate decreases further because mutations are too rare for even (1, 0) or (0, 1) to establish reliably. In this regime, the success rate is directly determined by the potential differences, because *P*_establishment_ ∝ |ΔΦ|, with ΔΦ_(1,0)←(0,0)_ = −0.5·*s*, ΔΦ_(0,1)←(0,0)_ = −0.8·*s*. Therefore, the success probability can be estimated as 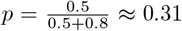 which is consistent with the value observed in Fig [5] for *m* ~ 10^−10^. It is important to note that variations in the selection coefficient *s* have only a minor effect: as inequality (5) shows, the parameter *α*, which primarily determines the probability of success, depends on the *proportion* of potential differences rather than their absolute values.

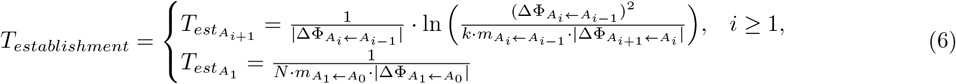

The theoretically estimated time for the integer evolutionary reaction to reach 1346 under the defined mutation scheme is calculated as *T* = 11451, assuming that when |*x* − 50 | ≤ 1346, there are fifteen +3 mutations and five +1 mutations. Depending on the assumption, the estimated time varies: *t* = 9044 for sixteen +3 and two +1 mutations, and *t* = 14782 for fourteen +3 and seven +1 mutations. This demonstrates that the expected time is path-dependent. The simulation result shows that the average time over 50 iterations is *t* = 11011 generations, suggesting that the dominant evolutionary path involves fifteen +3 and five +1 mutations, which is plausible given the preset mutation probabilities. The overall success rate from the simulation is 0.58, as the evolution may terminate at 648 due to early establishment of 3 and fixation of 18 before 6 is fixed.

For the modulo vector evolutionary reaction, the average times over 50 iterations are *t* = 3909 for the objective vector (1,1,1,1) and *t* = 15552 for the objective vector (2,3,1,2). These values differ by only 4% and 0.02% from the theoretical times *t* = 4070 and *t* = 15555, which are calculated by summing the establishment times from (6), a generalization of (4). In the first case, the paths are isotropic—potential changes are uniform regardless of which mutation occurs first—so the expected time does not depend on the specific path taken. In the second case, although the total time is path-dependent, the population tends to follow the path of maximal potential decrease, specifically:

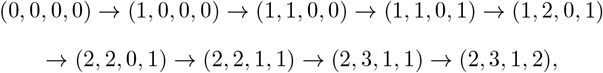

so that theoretical and simulation values are nearly identical.

The result for the 50 iterations of the 2 × 2 matrix demonstrates the expected convergent evolution property, producing the matrices

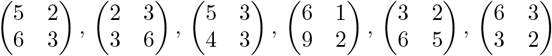

which indeed suffice the determinant 3 and trace 8 condition. Note that although a matrix such as 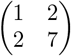 or its determinant and trace preserving permutation matrices also satisfy the potential minimum, these matrices were not observed in the simulation result.

Since many evolutionary paths can lead to the same resulting matrix, specifying the path is necessary to theoretically estimate the time of that path. However, comparing the measured and theoretical times for all possible paths is not done here, as the number of possible paths is large, and such a calculation is not essential.

For the case of cyclic algebra evolution, as illustrated by the Muller plots in Fig. 6, the equilibrium among the initial Rock, Paper, and Scissors states—each with equal proportion—is susceptible to stochastic disturbances, particularly when the selection coefficient is high. Furthermore, a higher mutation rate is observed to delay the disruption of this initial equilibrium while simultaneously promoting clonal interference. These simulation results indicate that cyclic evolution can indeed emerge from the defined potential.

**Figure 6.**
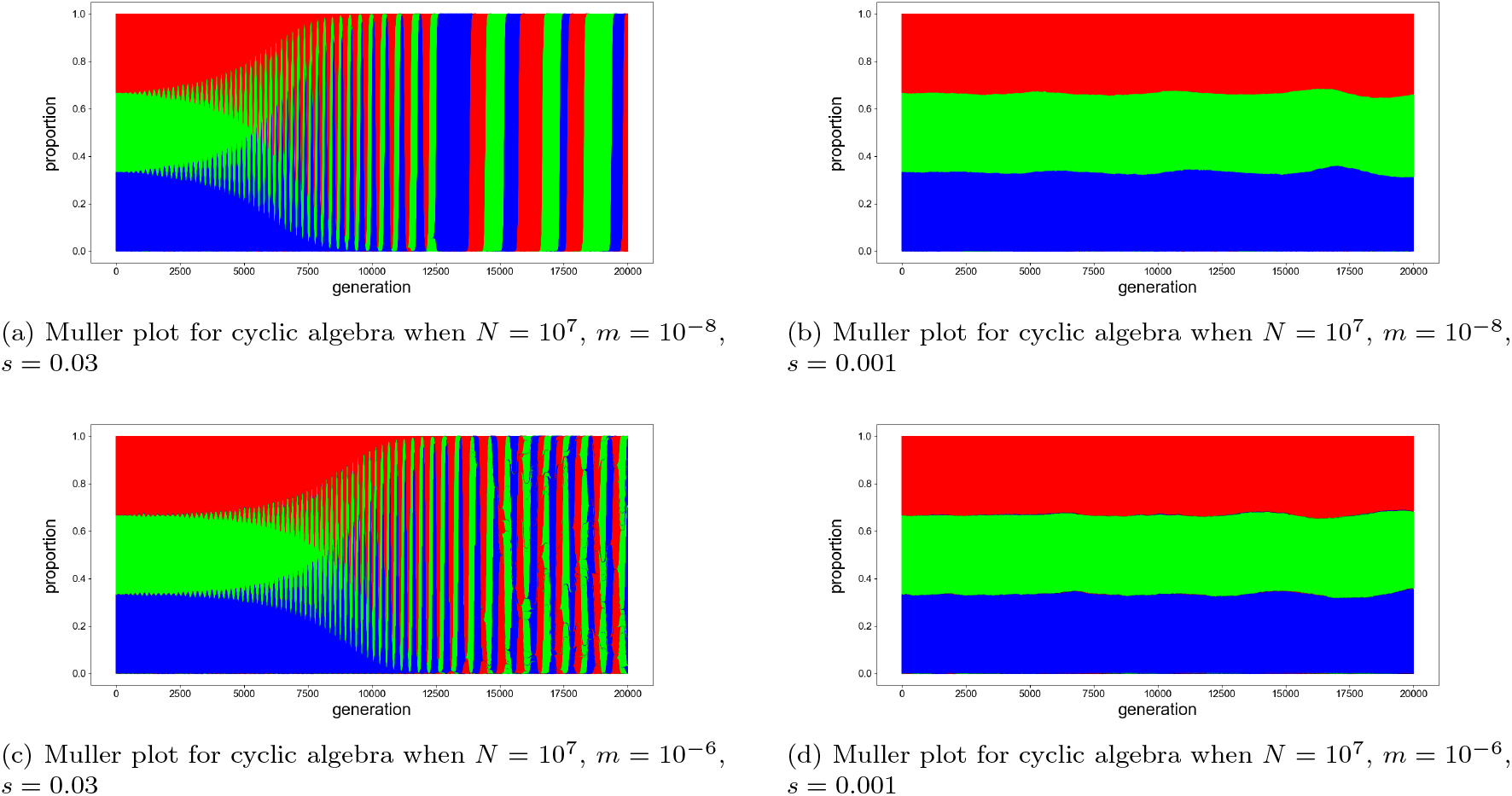
Muller plots for cyclic algebra with various evolutionary parameters.

The results demonstrate that general algebra can be utilized to model particular traits of interest in an asexual population. Moreover, when the potential and evolutionary pathway is specified, the deterministic behavior predicts both the eventual state of the algebra and the time required to reach it. While this phenomenon may appear similar to the physical potential that defines the state of an object in classical or quantum mechanics, there is a crucial difference. In physical systems, the potential can be independent of the particle’s state, whereas in evolutionary dynamics, the potential is fundamentally state-dependent: the current trait shapes the potential landscape. For example, in quantum mechanics, the wave function in an infinite potential well typically does not alter the potential as it evolves, but in evolutionary dynamics, the evolution of the state *melts* the initial potential, since the trait itself defines the potential’s shape. Therefore, ensuring conditions that allow an *effectively time-independent potential*, even as evolution proceeds, is essential for the methodological analysis described above.

The integer evolution case clearly demonstrates that, with a finite number of potentials, desired algebra can be evolved within the predicted timescale. It is important to note that the potential selected for the simulation represents only one of many possible choices; therefore, one may select a potential that maximizes the efficiency of DE. Moreover, if the potential decrease associated with powers of 6 is insufficient, fixation of unintended algebraic states may occur. To prevent this, one can either modify the potential or reduce the mutation rate for unwanted mutations during certain stages of evolution, albeit at the cost of slowing the whole evolutionary reaction. This integer-based algebra is applicable to traits that exhibit simple additive or multiplicative behavior across generations. For realistic applications, it is crucial to choose potentials that minimize evolution time and maximize the success rate, while taking into account constraints on the number of potentials.

The time required for the modulo element vector to reach the objective vector was consistent with the theoretically calculated timescale. This algebra is particularly useful for modeling the evolution of DNA fragments, where each number 0, 1, 2, 3 corresponds to one of the nucleotides A, G, T, and C. Naturally, such a vector representation can be extended to non-modulo vectors with different dimensions, as illustrated in the two-dimensional vector case above.

Next, the matrix evolution case demonstrates that the inner (or emergent) properties of the algebra determine its evolutionary dynamics and illustrate the occurrence of convergent evolution. For in-depth mathematical analysis, the local and global minimum points of the given potential can be calculated by taking partial derivatives with respect to the independent variables *a, b, c*, and *d*. The results show that the global minima lie on (*a* + *d*) = 8 and *ad* − *bc* = 3, with no local minima present. However, this does not ensure that the system can always reach the global minimum for set of mutation matrices.

For example, if the only allowed mutations are 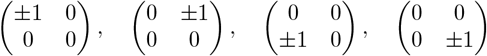 the matrix can get stuck in states such as 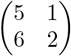 because no available mutation decreases the potential. Mutations like 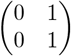 are therefore necessary to reach the objective, explaining why 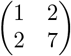 cannot be reached with the given mutation set.

Moreover, the example demonstrates that the predicted time for the evolutionary reaction depends on the chosen pathway. For example, if the matrix evolves along

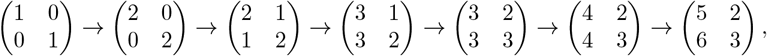

the total evolution time is *T* ≈ 30941, with the step 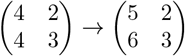 acting as a bottleneck. In contrast, following the path

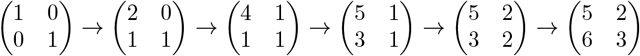

yields *T* ≈ 10047. This demonstrates that there exist various evolutionary paths along which the potential continuously decreases at every step until the algebra reaches the objective.

Here, we observe that when the decrease in potential is small—as in the bottleneck case—the establishment time is prolonged. Despite this, many evolutionary trajectories may fall into such traps because larger potential drops yield higher fixation probabilities. Consequently, evolution is more likely to proceed in the direction of −∇ Φ, favoring trajectories with large potential drops in the early stages. This bias leaves only small potential decreases for the later stages, which in turn prolongs the overall evolution time. Note that this behavior is observed in the 4-D modulo vector case for evolving (2,3,1,2).

Thus, the fastest evolutionary scenario occurs when the potential decreases nearly uniformly at each mutation step, while the total number of steps is minimized. Such fact can be formalized by simplifying Eq (6): 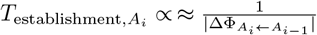, and applying Jensen’s inequality Jensen [1906] to the convex function:

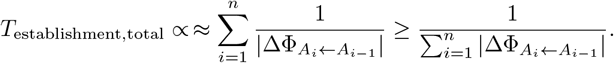

The minimum time is achieved when 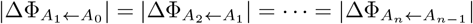. This principle applies to any algebra, suggesting that when the potential drop to the objective algebra is fixed, evolutionary pathway that offers nearly uniform gaps of potential drop raises the efficiency of the whole reaction.

Note that when designing such uniform gaps of potential is not feasible, increasing the mutation rate can activate relative tunneling and facilitate efficient evolution.

Another strategy to increase the efficiency of evolutionary reactions is to vary the mutation rates for specific traits, promoting beneficial evolutionary changes while limiting less advantageous ones. For instance, with 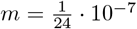 for all mutations, the path

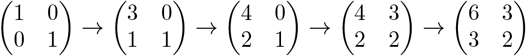

takes *T* = 4277. However, when mutation rates are adjusted to 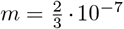 for + 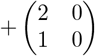 and 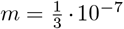 for 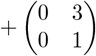, while other mutations are blocked, the time reduces to *T* = 3295.

Moreover, cyclic evolution demonstrates that evolution can eventually return to the same state. This phenomenon can be illustrated in a scenario where the mutation color of the prey is limited to R, G, and B, while the predator’s color recognizability co-evolves to better detect the color of the prey that is more abundant in the population. In nature, since evolution is based on DNA mutations, and the number of base pairs in bacteria is on the order of 10^5^ ~ 10^7^ Land et al. [2015], cyclic evolution at the DNA level would be statistically difficult to observe unless specialized methodologies are employed to selectively mutate particular sequences Park and Kim [2021], Chen et al. [2025] *in vivo*. However, at the level of general phenotypical traits, when these traits are well-defined and categorized, cyclic evolution could still be observed even without specialized molecular biological interventions.

The key assumptions for the algebraic modeling are: (i) The algebra represents phenotypical traits in a general sense, which can include genetic codes whenever the latter are interpreted as phenotypical traits; (ii) Mutations in the algebra are defined at the level of phenotypical traits and act in a manner that ensures the set of algebraic elements remains closed; and (iii) The potential of a population state can be considered effectively time-independent. To implement these assumptions in the laboratory, several considerations are necessary. First, there should be well-defined criteria linking phenotypical traits to specific algebraic representations. It should be noted that *m*_Δ**x**+**x**←**x**_ may be difficult to predict if the categorization allows excessive variance in the underlying genetic information. Therefore, to achieve a well-defined categorization while restricting the range of possible genetic information corresponding to a trait, one should design an algebra with sufficient sophistication. That is, although the algebra need not encode every detail of the genetic information in a one-to-one manner, it should capture enough subtleties of the trait to eliminate unwanted ambiguities and large variances. When the algebra is well defined, it would also become possible to investigate methods for controlling DNA mutations that correspond to algebraic mutations.

Second, to ensure that the algebraic set remains closed, the simplest approach is to disregard mutations that fall outside the defined algebra. Moreover, by employing biological techniques that enable *in vivo* selective mutations at specific regions of DNA Park and Kim [2021], Chen et al. [2025], it is possible to restrict mutations to occur only within the algebraic set.

Third, to maintain a locally time-independent potential, the structure of the potential Φ = Φ(*A, A*^*C*^) should be taken into account. In this formulation, the traits corresponding to the algebra set *A* of interest, as well as other traits represented by *A*^*C*^, all contribute to the overall potential. With such a structure, the time dependence of the potential may arise either from the evolution of the algebra *A* itself or from the evolution of the complementary algebras *A*^*C*^. In the former case, one way to maintain an effectively fixed potential is to consider short generational intervals and recalculate the potential as the state moves along its evolutionary path. When the time dependence originates from the evolution of *A*^*C*^, in addition to recalculating the potential, the mutation rate can be selectively increased for genes corresponding to *A* using *in vivo* biotechnological methods. This strategy makes the evolution of *A*^*C*^ relatively slower than that of *A*, thereby ensuring that the potential remains effectively time-independent over a certain range of generations.

Although the aim of this paper was to examine the general behavior of algebraic evolution and to demonstrate that such modeling is feasible, defining a specific algebra that accurately represents a particular trait remains an important task. One of the general approaches would be to use an *n*-dimensional manifold *M* to model *n* independent traits of interest and perform algebraic analysis on some smooth function *f* such that *f* ∈ *C*(*M*) = { *f*: *M* → ℝ^*n*^}, drawing from the framework of algebraic geometry Hartshorne [2013]. Note that such a manifold can be defined in any abstract *n*-dimensional space spanned by basis elements derived from the *n* traits. Moreover, we could approximately represent such *f* (*M*) with a *N*_1_ × *N*_2_ ×…*N*_*n*_ matrix by *discretization* Krispel et al. [2015] of such continuous geometry while in this case, the element of the matrix mutates. However, constructing such a manifold—or devising other general algebras that appropriately represent the traits of interest—extends beyond the scope of this paper and requires further study. The essential point is that, regardless of the specific algebraic structure chosen to represent a trait, that structure will always follow the deterministic behavior governed by the definitions and conditions introduced above.

Finally, to rigorously predict the success rate of the DE reaction within a given range of generation intervals, it is essential to determine the distribution of the establishment time that corresponds to the general algebraic case in inequality (5). If such a distribution is determined, the probabilities corresponding to a given generation interval for each step along the evolutionary path can be calculated rigorously. These probabilities can then be combined—through multiplication and summation—to obtain the overall success rate of the evolutionary reaction. Therefore, further research aimed at determining the distribution of the establishment time is of significant importance. significant importance.

## 5 Conclusion

The algebraic model captures the intuition underlying phenotypical trait-based DE, which humans have been successfully practicing for thousands of years. With this promising framework, the directed evolution (DE) of asexual populations can be quantitatively analyzed. Specifically, once an algebra is well-defined to represent a particular trait, the evolutionary pathway can be designed using the corresponding potential and mutation rates, allowing the relevant evolutionary quantities to be calculated. Simulation results for diverse example algebras matched the theoretical predictions for the number of generations required, demonstrating that a quantitative analysis of directed evolution is indeed feasible. Finally, possible improvements to the algebraic model were also discussed. Although further development is needed, once fully established, the model is expected to provide a robust framework for artificially guiding the evolution of asexual populations in a controlled and effective manner.

## 6 Data Availability

The code for the simulation could be found from https://github.com/TheDEnotes/Algebraic-modeling-for-Directed-Evolution.

## 7 Acknowledgement

This work was developed mainly at the Department of Physics, Yonsei University, and at DAMTP, University of Cambridge. I sincerely appreciate the calm and supportive environments offered by both institutions.

## 8 Study Funding

This is a self-funded research.

## 9 Conflict of Interest

There is no conflict of interest to declare.

## Appendix

Fig. A[1] demonstrates the potential for integer evolution. The red dots and line indicate the expected trajectory of the integers throughout the evolutionary process.

**Figure A1.**
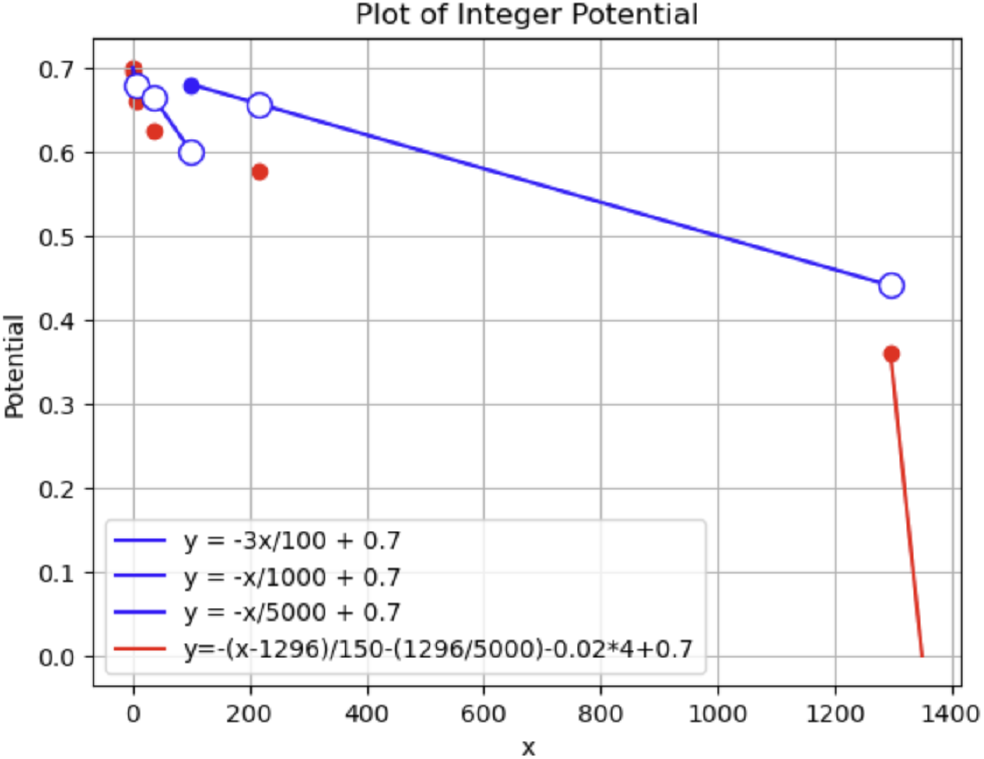
Potential for the integer evolution

The mutation matrices used for the above example are the following

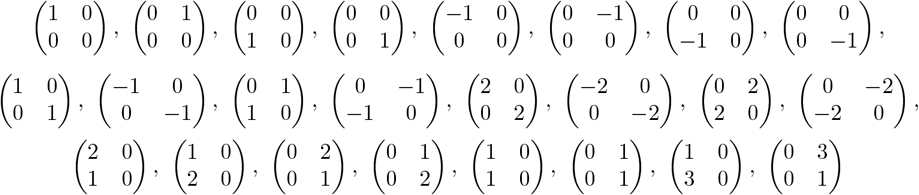

with all having a mutation rate of 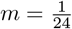.

## Notes

https://github.com/TheDEnotes/Algebraic-modeling-for-Directed-Evolution

### Competing Interest Statement

The authors have declared no competing interest.

### Summary of Updates

Minor modifications for the representations are made.

